# Robust Prediction of Patient-Specific Cancer Hallmarks Using Neural Multi-Task Learning: a model development and validation study

**DOI:** 10.1101/2025.02.03.636380

**Authors:** Shreyansh Priyadarshi, Camellia Mazumder, Bhavesh Neekhra, Sayan Biswas, Debojyoti Chowdhury, Debayan Gupta, Shubhasis Haldar

## Abstract

**Background:** Accurate quantification of cancer hallmark activity is essential for understanding tumor progression, tailoring treatments, and improving patient outcomes. Traditional methods, such as histopathological grading and immunohistochemistry for protein expression, often overlook the complex interplay between cancer cells and the tumor microenvironment and provide limited insight into hallmark-specific mechanisms. We aimed to develop OncoMark, a high-throughput deep learning-enabled neural multi-task learning framework capable of systematically quantifying integrative hallmarks activities using transcriptomics data from routine tumor biopsies.

**Methods:** In this study, we acquired single-cell transcriptomics data from 941 tumor samples across 14 tissue types, comprising nearly 3.1 million cells from 56 studies conducted worldwide, to form a large multicenter dataset. Our model employs a supervised neural multi-task learning method designed to predict multiple cancer hallmarks present in the biopsy samples simultaneously. The OncoMark model was developed and tested on 90% of the studies (patients from 51 studies) using repeated five-fold cross-validation performed twice. For further evaluation, the model was assessed on the remaining 10% of the studies (patients from 5 studies) that were excluded from the initial training and testing dataset. Additionally, we included patients from publicly available datasets, including TCGA, GTEx, ANTE, MET500, POG570, CCLE, TARGET, and PCAWG to validate the model’s performance. The primary objective was to evaluate the performance of the model in identifying cancer hallmarks in cancer datasets and ensure no hallmark predictions were made in normal samples across the four prespecified groups: (i) internal test set, (ii) external test set, (iii) normal samples (real-world), and (iv) cancer samples (real-world).

**Findings:** OncoMark demonstrated exceptional performance in predicting cancer hallmark states, achieving near-perfect accuracy across internal test data and five external test datasets. Internal testing consistently showed accuracy, precision, recall, and F1 scores exceeding 99%, underscoring the model’s reliability across hallmarks. External test further confirmed these findings, with accuracy, precision, recall, F1 scores, and balanced accuracy consistently exceeding 96·6%, and multiple datasets achieving perfect scores, highlighting the model’s exceptional generalizability and robustness. Specificity tests using GTEx and ANTE datasets accurately classified normal tissues, while sensitivity analysis on TCGA, MET500, CCLE, TARGET, PCAWG, and POG570 datasets effectively identified cancer hallmarks.

**Interpretation:** We developed an AI-based framework that enables accurate, efficient, and cost-effective quantification of cancer hallmark activity directly from transcriptomics data. The framework demonstrated significant potential as an assistive tool for guiding personalized treatment strategies and advancing the clinical management of cancer patients.

**Funding:** Ashoka University, S.N. Bose National Centre for Basic Sciences, Mphasis F1 Foundation, DST SERB Core Research Grant.

**Research in Context:** *Evidence before this study:* We conducted an extensive literature search using Google Scholar and PubMed without language restrictions, employing search terms such as “(Predicting OR Classifying OR Annotating) and (cancer hallmarks) AND (Deep OR Machine Learning) OR (Artificial Intelligence OR AI).” While there have been advancements in molecular oncology and computational methodologies over the two decades since the concept of cancer hallmarks was first introduced, a comprehensive machine learning or deep learning framework to annotate all cancer hallmarks simultaneously from tumor biopsy samples remains to be developed. Additionally, the scarcity of hallmark-annotated datasets has posed a significant challenge, hindering the development of robust predictive models.

*Added value of this study:* This study introduces OncoMark, a novel high-throughput neural multi-task learning (N-MTL) framework designed to predict all cancer hallmark activities simultaneously from biopsy samples. OncoMark addresses the lack of annotated hallmark-specific data by generating synthetic biopsy (pseudo-bulk) datasets annotated with hallmark activity, meticulously modeled to reflect real-world tumor biology while maintaining clinical relevance. The framework employs a multi-task learning approach to capture interdependencies among hallmarks, advancing beyond isolated predictions to offer a holistic view of tumor biology. Validation on five independent datasets comprising 95 patient samples demonstrated its generalizability and reproducibility. Further external validation using eight datasets, encompassing over 11,679 cancer and 8348 normal patient samples, reinforced its robustness. To promote clinical integration, a user-friendly web-based tool was developed, enabling seamless access for oncologists and researchers.

*Implications of all the available evidence:* The OncoMark framework represents a transformative advancement in cancer diagnostics and treatment planning. By enabling accurate and reproducible prediction of all hallmark activities simultaneously from biopsy samples, this model paves the way for precision oncology at scale. Its ability to systematically capture hallmark interdependencies provides deeper insights into tumor behavior, guiding the development of individualized targeted therapies. The incorporation of a web-based interface ensures the accessibility of this innovation to clinicians worldwide, bridging the gap between computational oncology and clinical practice. Following further validation and integration into healthcare workflows, OncoMark has the potential to improve cancer outcomes by delivering timely, cost-effective, and precise tumor analyses, facilitating informed therapeutic decision-making with unparalleled precision.

Cancer progression is driven by a set of well-defined biological principles—collectively termed the “hallmarks of cancer”—yet current diagnostic approaches seldom incorporate these distinct molecular features into clinical practice. Despite substantial progress in molecular oncology, traditional methods like histopathological grading and immunohistochemical assays often fail to capture the complex interplay between cancer cells and the tumor microenvironment, emphasizing the need for robust computational frameworks capable of systematically quantifying hallmark-specific activity. Here, we address this gap by developing OncoMark, a high-throughput neural multi-task learning (N-MTL) framework designed to simultaneously quantify hallmark activities in tumor biopsies using transcriptomics data. We show that OncoMark achieves near-perfect accuracy, precision, recall, and F1 scores (>99%) in cross-validation, with external validation consistently exceeding 96.6% on five independent datasets. Further evaluation on eight additional datasets—including large-scale cancer cohorts (TCGA, MET500, CCLE, TARGET, PCAWG, POG570) and normal tissue datasets (GTEx, ANTE)—demonstrated high specificity for normal samples and robust sensitivity for hallmark prediction in cancer. By delivering a comprehensive and cost-effective molecular portrait of tumor biology and providing a user-friendly web platform accessible at https://oncomark-ai.hf.space/, OncoMark has the potential to guide tailored treatment strategies and advance precision oncology. More broadly, this framework signifies a transformative step toward routine hallmark-based diagnostics, promising to improve patient outcomes by facilitating timely and precise tumor profiling.

## Introduction

Cancer is an inherently heterogeneous disease, yet it progresses through well-defined biological principles that govern its development and spread.^1,2^ Hanahan and Weinberg’s introduced the concept of the “hallmarks of cancer,” a unifying framework that identifies the fundamental capabilities cancer cells acquire during tumorigenesis.^3^ These core hallmarks include: (1) sustaining proliferative signaling to drive uncontrolled growth; (2) evading growth suppressors to bypass regulatory constraints; (3) resisting cell death to survive environmental and intracellular stress; (4) enabling replicative immortality to achieve limitless cell division; (5) inducing angiogenesis to ensure a continuous nutrient supply through neovascularization; and (6) activating invasion and metastasis to colonize distant tissues. This framework has been expanded to incorporate emerging hallmarks, such as (7) deregulating cellular energetics to sustain rapid proliferation, and (8) avoiding immune destruction by escaping immune surveillance. Enabling characteristics, including (9) genome instability and mutation, which accelerate tumor evolution, and (10) tumor-promoting inflammation, which supports a microenvironment conducive to malignancy, further illustrate the complexity of cancer biology.^4^

Despite the insights offered by these hallmark framework, current diagnostic approaches often fail to integrate these molecular underpinnings into routine clinical practice. Traditional methods, such as staging systems (e.g., AJCC, TNM) and grading scales (e.g., Gleason grading), primarily focus on macroscopic and microscopic tumor characteristics, overlooking the molecular heterogeneity that drives tumor behavior.^5,6^ Consequently, patients with the same cancer type, stage, and grade may exhibit divergent outcomes, exposing the limitations of these approaches.

Moreover, these approaches do not provide insights into the dynamic, micro-evolutionary molecular changes within tumors, limiting their capacity to guide personalized treatment strategies.^7^ A hallmark-based diagnostic framework has the potential to address these limitations by integrating molecular data to illuminate the biological mechanisms underlying tumorigenesis— an essential step toward precision oncology, in which treatments are tailored to the individual tumor’s molecular profile.^8^ Although multi-omics technologies, artificial intelligence, and real-time monitoring have advanced considerably, we still lack a single, unified method that can simultaneously annotate all hallmark activities in a tumor.^9,10^.

To address this critical gap, we developed OncoMark, a high-throughput neural multi-task learning (N-MTL) framework designed to simultaneously quantify the activity of all cancer hallmarks using transcriptomic data from tumor biopsies. This is, to the best of our knowledge, the first computational tool specifically designed to predict all cancer hallmarks concurrently. Upon input, OncoMark calculates the probability of each hallmark’s activity, providing a detailed molecular profile of the tumor. The model underwent a rigorous validation process to ensure robustness and generalizability. Cross-validation demonstrated accuracy, precision, recall, and F1 scores exceeding 99%. Testing on five independent external datasets^11,13,14,15,16^ further confirmed its performance, consistently maintaining a minimum of 96.6% across these metrics. Additional validation on eight gold-standard datasets, comprising six cancer (TCGA, MET500, CCLE, TARGET, PCAWG, and POG570) and two normal (GTEx and ANTE) datasets, verified the model’s high accuracy in distinguishing true positives and true negatives. Moreover, predicted hallmark activity demonstrated significant co-association with AJCC stages and TNM staging, with the strongest co-association observed at advanced stages of cancer progression. Building on its clinical utility, we have developed a user-friendly software platform, accessible at https://oncomark-ai.hf.space/, which seamlessly integrates hallmark activity profiling into research and clinical workflows.

## Methods

### Study Design

In this study, we developed OncoMark, a novel neural multi-task learning framework designed to predict cancer hallmark activity from transcriptomic data obtained from biopsy samples (figure 1). Direct experimental methods for annotating biopsies transcriptomics with hallmark activity are yet to be established, and as a result, no hallmark-annotated biopsy datasets were available for model training. To overcome this limitation, we generated synthetic bulk biopsy data utilizing single-cell transcriptomes to emulate real tumor conditions. We manually curated and compiled gene sets associated with each hallmark. These gene sets were used to calculate digital scores for each single cell, reflecting hallmark activity. The digital scores were subsequently binarized to produce binary annotations indicating the presence or absence of each hallmark in individual cells. Synthetic biopsy data, annotated for hallmark presence or absence, were then generated by leveraging cells exclusively from each biopsy sample. This approach faithfully emulates real-world tumor conditions while preserving biological integrity and ensuring no cross-sample contamination. Moreover, cancer hallmarks do not function independently; they interact and influence one another. Capturing these interdependencies is critical for building accurate predictive models. To address this, we avoided training models on individual hallmark datasets in isolation. Instead, we employed a neural multi-task learning framework that first learns shared representations across all hallmark datasets. Subsequently, hallmark-specific neural networks are trained using this shared representation, enabling the model to effectively capture hallmark interdependencies. The model was rigorously validated using intrinsic test datasets, external test datasets, and real-world tumor and normal datasets, ensuring its robustness and generalizability across diverse conditions.

**Figure 1:**
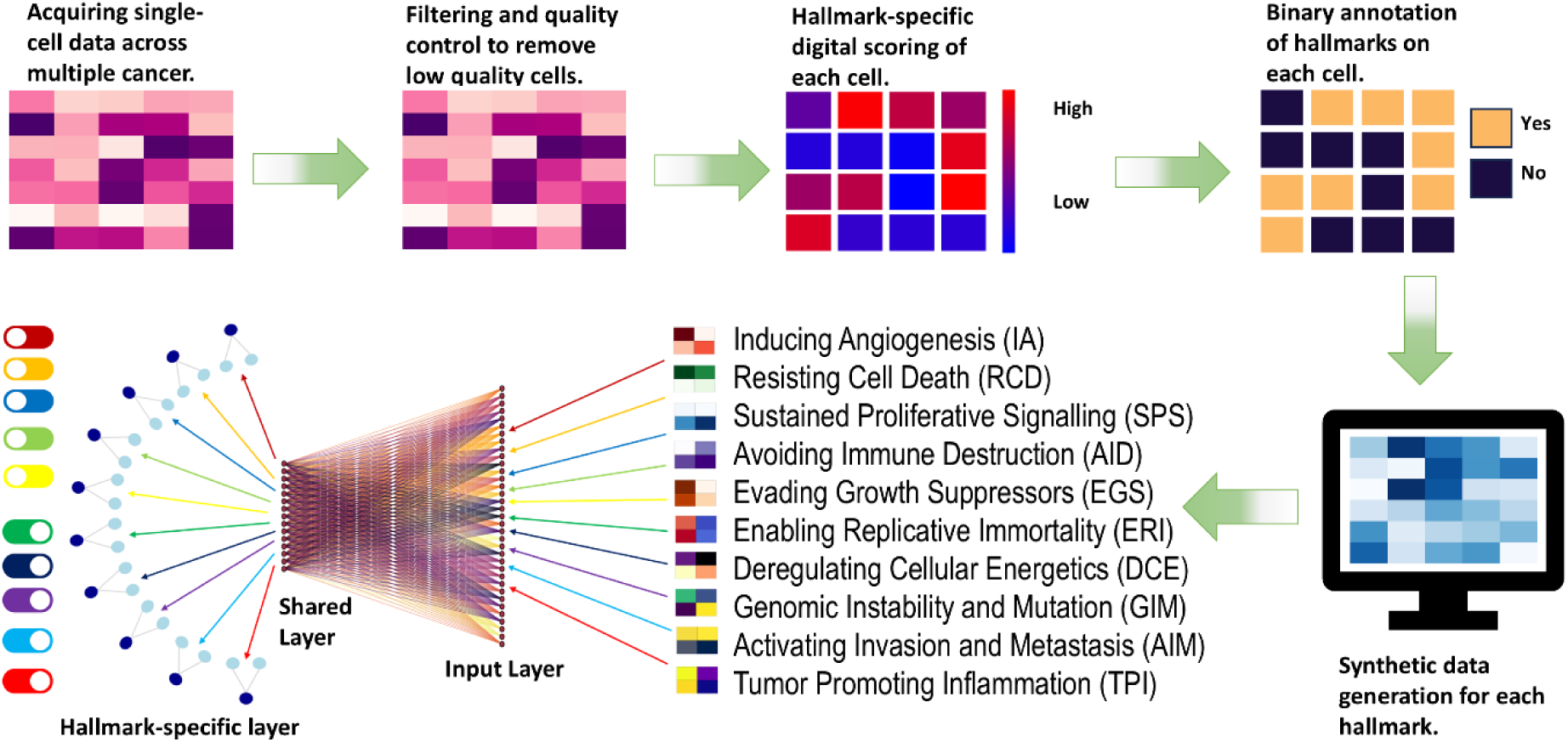
Workflow for Cancer Hallmark Prediction Using OncoMark. Overview of the pipeline for predicting cancer hallmarks using OncoMark. Single-cell data from multiple cancer types undergoes quality control to filter out low-quality cells. Hallmark-specific gene expression is digitally scored, producing binary annotations (Yes/No) for hallmark presence. Annotated cells are used to generate synthetic (pseudo-bulk) datasets for each hallmark, which are then utilized to train a Multi-Task Neural Network (M-TNN). The M-TNN learns a shared representation across all hallmarks, followed by hallmark-specific layers for precise hallmark prediction.

Beyond its scientific contributions, OncoMark was developed with practical application in mind. A web-based tool was created to enable clinicians to upload tumor transcriptomics data and receive real-time predictions of hallmark activity. This tool might provide actionable insights to support clinical decision-making and personalize treatment strategies, considering the unique biological characteristics of each tumor.

### Data Preparation

We used a comprehensive dataset of 3.1 million single-cell transcriptomes from 14 tumor types, collected from 941 patients across 56 studies worldwide as part of the Weizmann 3CA repository, to generate synthetic (pseudo-bulk) datasets. For model training and cross-validation, we used 90% of the studies (encompassing 846 patients from 51 studies). For external validation, we used the remaining 10% of studies, which were excluded from the training set. These five excluded studies — Dong et al. (2020),^11^ Yost et al. (2019),^12^ Pal et al. (2021),^13^ Gao et al. (2021),^14^ and Nam et al. (2019)^15^ — included a total of 95 patients. Additionally, we collected real-world bulk transcriptomic data from publicly available sources, including The Cancer Genome Atlas (*TCGA*, n=6679), MET500 (n=868),^16^ POG570 (n=570),^17^ Cancer Cell Line Encyclopedia (*CCLE, n=1019*),^18^ Therapeutically Applicable Research to Generate Effective Treatments (*TARGET, n=734*), Pan-Cancer Analysis of Whole Genomes (*PCAWG, n=1521*),^19^ and normal datasets from Genotype-Tissue Expression (*GTEx, n=8228*),^20^ and Atlas of Normal Tissue Expression (*ANTE, n=120*).^21^ These datasets were used to evaluate the sensitivity and specificity of *OncoMark* in predicting hallmark activity in both synthetic and real-world settings. All datasets were publicly available, de-identified, and obtained with appropriate consent from participants in their respective studies. Institutional permissions for data use were secured by respective studies, negating the need for further ethics approvals.

Before synthetic data generation, single-cell data underwent a stringent quality control process. Cells with mitochondrial transcript content exceeding 15% or expressing fewer than 200 or more than 6,000 mRNA transcripts were excluded.^22,23^ Gene sets associated with cancer hallmarks were curated from multiple databases, incorporating only genes identified in at least two independent sources.^24–29^ The hazard ratio of each gene was calculated using the Cox proportional hazards model on the TCGA dataset.^30^ Genes with hazard ratios below 1.05 were excluded from further analysis. The initial selection was further refined by a manual review of the literature to check for genes directly or indirectly involved in hallmark-related pathways. Using these curated gene sets (table S1 appendix 1 pp 15-23), we computed Digital Scores for each of the 10 cancer hallmarks across all 2·7 million cells, employing the UCell.^31^ Hallmark presence or absence was determined via Otsu’s thresholding method, ensuring robust binary classification. Given the tissue-specific nature of hallmark expression, we calculated tissue-specific digital score thresholds to distinguish hallmark-positive and hallmark-negative cells for each hallmark.

To simulate real-world biopsy conditions while preserving biological fidelity, we aggregated 200 hallmark-specific cells from each patient sample to create synthetic biopsy datasets.^32,33^ Cells were grouped by hallmark status (positive or negative) to generate hallmark-specific synthetic biopsies for each status, ensuring no overlap across synthetic samples and minimizing cross-sample contamination. Separate synthetic datasets with positive and negative ground truths were generated for each hallmark to facilitate robust model training and to mimic the heterogeneous composition of clinical samples. For validation on five external studies, the synthetic data creation steps applied to training data were replicated. However, for each patient, hallmark-positive cells were pooled to create positive datasets, and hallmark-negative cells were pooled to create negative datasets, resulting in data that more closely resembles real-world bulk RNA sequencing profiles. This approach ensured consistency between training and validation while enabling the model to generalize to clinically relevant bulk transcriptomic datasets.

To mitigate batch effects and reduce overfitting risks, both synthetic biopsy and real-world biopsy datasets were transformed into rank space.^34^ The data were subsequently log2-transformed and standardized such that each feature had a mean of zero and a standard deviation of one.^35^ These preprocessing steps facilitated faster convergence during model training and improved the interpretability of the learned features.

### OncoMark Framework

The OncoMark Framework leverages deep learning approach to predict hallmark activity in biopsy samples, emphasizing both hallmark-specific precision and the biological interplay among hallmarks. The model architecture is structured as a multi-task neural network, consisting of a shared base layer and task-specific output layers (figure S3 appendix 1 p 10).^36^ The shared base layer processes the input features to extract pan-hallmark characteristics that are universal across all cancer hallmarks. The task-specific output layers then refine these shared representations by focusing on hallmark-specific features, enabling the model to capture the nuanced interplay between hallmarks. This architectural design reflects the interconnected nature of hallmark activities observed in tumor biology and ensures that predictions are biologically meaningful. By combining a shared representation with hallmark-specific refinement, the framework achieves accurate predictions that align with the cooperative and dynamic behavior of hallmarks in cancer progression. The model outputs independent probability scores for ten hallmarks simultaneously, which can be interpreted as the probability of hallmark presence. The detailed model’s architecture, and training methodology are elaborated in the supplementary appendix (appendix 1 pp 5-10).

The model was trained on a balanced dataset comprising 67,930 samples with 9,326 input features representing gene expression profiles. Of these, 57,735 samples (85%) were used for training, and 10,195 samples (15%) were used for validation, with data splitting performed separately for each hallmark using the train-test split method to ensure balanced representation across both sets. To prevent catastrophic forgetting, data from all hallmark tasks were merged and shuffled randomly during training, avoiding hallmark-specific batches and ensuring uniform exposure of the model to all hallmark datasets. The Adam optimizer, with a learning rate of 0·0001, was employed to minimize the binary cross-entropy loss, which was calculated independently for each hallmark prediction task and combined into a weighted average using task-specific indicators to ensure balanced learning. Early stopping, with a patience of six epochs, was implemented to prevent overfitting by halting training when validation loss showed no improvement, while a learning rate scheduler further adjusted the learning rate by reducing it by 0·5 after three consecutive stagnant epochs, with a minimum threshold set at 1e-6. Training was conducted for 50 epochs with a batch size of 256. Although validation losses consistently improved, gains became marginal in later epochs, leading to the decision to halt training after 50 epochs before full convergence (figure S4 appendix 1 pp 11, table S3 appendix 1 pp 23-25). To evaluate the model’s robustness and check for overfitting or data leakage, it was tested on external datasets comprising 95 tumor samples from five independent studies, with performance metrics such as F1-score, accuracy, precision, recall, balanced accuracy, precision-recall curve, and sensitivity-specificity curve confirming its ability to generalize effectively across diverse datasets.

### Statistical analysis

We employed a rigorous statistical framework to evaluate the performance of OncoMark. Our methodology incorporated five-fold cross-validation repeated twice to ensure robust model assessment (table S2 appendix 1 p 23). In each iteration, four folds were used to train the model, while the fifth fold was divided into two equal parts: one part served as a validation set for model selection and performance monitoring, and the other part was reserved for independent performance evaluation. Model performance was assessed using the F1-score, accuracy score, precision score, recall score, balanced accuracy, confusion matrix, area under the precision-recall curve (AUC-PR) and the receiver operating characteristic curve (AUC-ROC) (figure S5 appendix 1 pp 12, table S4 appendix 1 pp 25-34). We report the mean and standard deviation (SD) of these AUC values across all repetitions of cross-validation to capture variability and reliability. Additionally, the model was applied to two normal datasets, Genotype-Tissue Expression (GTEx) and Atlas of Normal Tissue Expression (ANTE), as well as six cancer datasets from The Cancer Genome Atlas (TCGA), MET500, POG570, Cancer Cell Line Encyclopedia (CCLE), Therapeutically Applicable Research to Generate Effective Treatments (TARGET), and Pan-Cancer Analysis of Whole Genomes (PCAWG) (figure S6 appendix 1 pp 13-14). Probability density distributions of hallmark predictions were plotted to assess the model’s ability to identify hallmark occurrences in cancer datasets, with no hallmark predictions made for normal samples. To determine whether the probability distributions between cancer and normal samples differ significantly, the Kolmogorov-Smirnov (K-S) test was conducted, providing statistical insights into these differences.^37,38^

Given the critical role of clinical staging in hallmark progression, we extended the model’s application to The Cancer Genome Atlas (TCGA) data to further investigate hallmark co-occurrence patterns across various AJCC and TNM staging systems. We quantified co-occurrence using odds ratios (ORs) to assess the strength of associations between specific hallmarks and their corresponding clinical stages.^39^ The results were visualized in heatmaps, with significant hallmark associations indicated by asterisks, based on a predefined statistical threshold of p-value < 0·05.

Furthermore, we examined the impact of cancer therapies on patient outcomes, specifically Overall Survival (OS), Disease-Free Survival (DFS), and Progression-Free Survival (PFS). Using TCGA data, we investigated how cancer drugs influence hallmark activity. Associations between hallmarks and drug treatments were quantified for all cancer patients and incorporated as features in a logistic regression model, with survival status as the dependent variable. The feature weights derived from the trained logistic regression model were interpreted as impact scores, offering a quantitative measure of each drug’s contribution to improving patient survival. These impact scores were used to rank the efficacy of different treatments in terms of their ability to alter hallmark pathways.

## Results

We evaluated our model performance in predicting cancer hallmarks during five-fold cross validation repeated twice. Our model performance in predicting AIM hallmark achieved an accuracy of 99·97% and an F1 score of 99·97%, with a balanced accuracy of 99·97%, tested on 729 patients (3364 positive and 4184 negative samples). Similarly, the DCE hallmark demonstrated an accuracy of 98·91% and an F1 score of 97·99%, achieving a balanced accuracy of 98·88% across 764 patients (3841 positive and 4249 negative samples). The EGS hallmark delivered exceptional performance with an accuracy of 99·95% and an F1 score of 99·92%, supported by a balanced accuracy of 99·96%, evaluated on 711 patients (3724 positive and 3168 negative samples). The GIM hallmark attained an accuracy of 98·73% and an F1 score of 98·68%, with a balanced accuracy of 98·74% across 395 patients (1282 positive and 846 negative samples). The RCD hallmark showcased strong metrics, achieving an accuracy of 99·93%, an F1 score of 99·92%, and a balanced accuracy of 99·93%, tested on 652 patients (2590 positive and 2833 negative samples). The SPS hallmark exhibited perfect performance, achieving 100% accuracy, F1 score, and balanced accuracy, validated on 695 patients (3305 positive and 3796 negative samples). The AID hallmark achieved an accuracy of 99·71% and an F1 score of 99·96%, with a balanced accuracy of 99·74%, evaluated across 691 patients (2832 positive and 3671 negative samples). The IA hallmark recorded an accuracy of 99·92% and an F1 score of 99·87%, with a balanced accuracy of 99·91%, tested on 705 patients (2318 positive and 4334 negative samples). The ERI hallmark demonstrated an accuracy of 99·28% and an F1 score of 98·53%, alongside a balanced accuracy of 99·08%, across 816 patients (3205 positive and 7238 negative samples). Lastly, the TPI hallmark achieved an accuracy of 99·43% and an F1 score of 99·13%, with a balanced accuracy of 99·57%, tested on 730 patients (2326 positive and 4824 negative samples) (table 1). For the ROC curves, all cancer hallmark predictions achieved near-perfect True Positive Rates (TPR) across all thresholds, with AUROC values of 1·00 ± negligible standard deviations, indicating robust discrimination between positive and negative samples for each hallmark (figure 2A). Similarly, the precision-recall curves confirm the models’ ability to maintain high precision at varying recall levels for each hallmark. The precision values remained consistently high, even as recall approached 1·0, highlighting the reliability of predictions in identifying positive cases without sacrificing precision (figure 2B).

**Figure 2:**
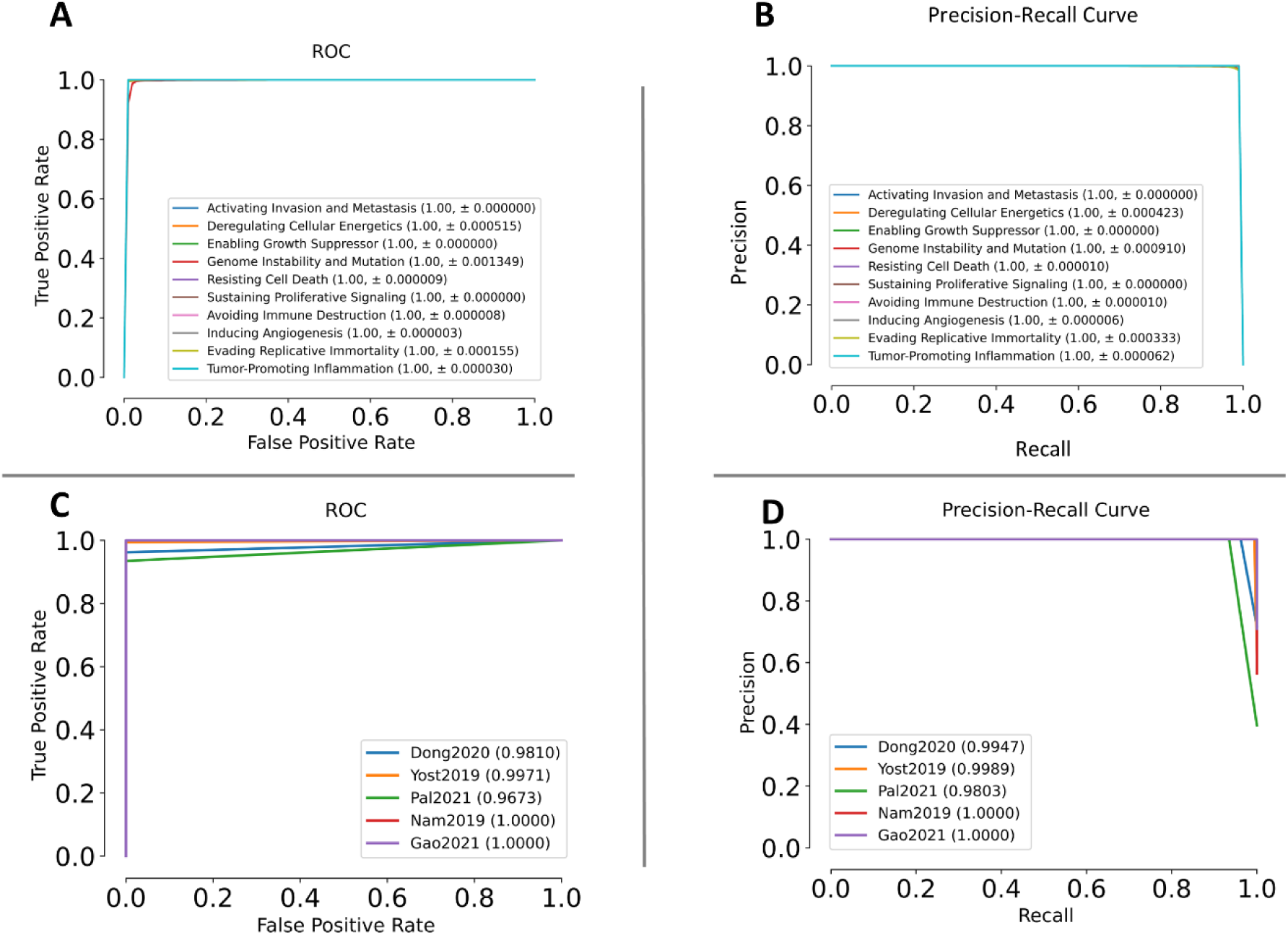
Evaluation of Hallmark Predictor Performance on Test and External Validation Datasets. (A) Mean ROC Curve and (B) Mean Precision-Recall Curve on Intrinsic Test Datasets from 5-fold cross-validation repeated twice. (C) ROC Curve and (D) Precision-Recall Curve on External Validation Datasets across all hallmarks. Labels indicate each hallmark along with the Area Under the Curve (AUC) ± standard deviation.

**Table 1:**
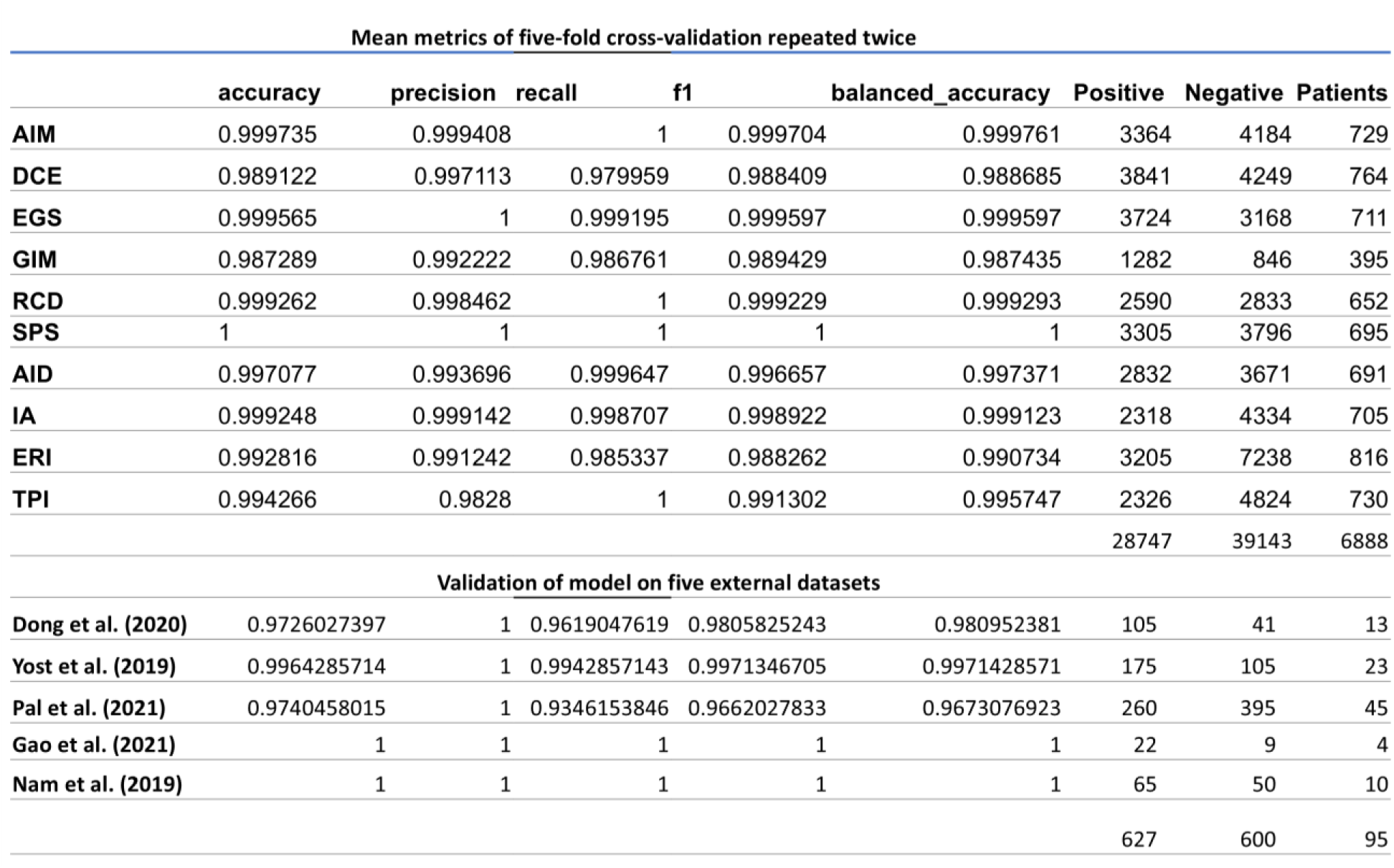
Sample information along with performance evaluation of OncoMark for the prediction of 10 cancer hallmarks. The model’s predictive accuracy was assessed using 5-fold cross-validation repeated twice on the primary dataset and validated on five independent external datasets. Metrics include accuracy score, precision score, recall score, f1-score, and balanced accuracy. Sample information includes the number of positive and negative samples along with the number of patients from which the given samples were generated.

The models were further validated on five external datasets. On the dataset from Dong et al. (2020), an accuracy of 97·26% and an F1 score of 96·19% were achieved across 13 patients (105 positive and 41 negative samples). Yost et al. (2019) reported an accuracy of 99·64% and an F1 score of 99·42% on 23 patients (175 positive and 105 negative samples). The dataset from Pal et al. (2021) yielded an accuracy of 97·40% and an F1 score of 93·46%, validated on 45 patients (260 positive and 395 negative samples). Both Gao et al. (2021) and Nam et al. (2019) achieved perfect metrics with 100% accuracy and F1 scores on 4 patients (1 positive and 22 negative samples) and 10 patients (65 positive and 50 negative samples), respectively (table 1). The ROC curves show high AUC values for datasets from Dong et al. (2020) (0·98), Yost et al. (2019) (0·99), and Pal et al. (2021) (0·97), whereas datasets from Gao et al. (2021) and Nam et al. (2019) achieved perfect AUC scores of 1.0 (figure 2C). Similarly, precision values remained close to 1·0 across recall levels for most datasets, with Dong et al. (2020) (0·99), Yost et al. (2019) (0·99), and Pal et al. (2021) (0·98) achieving excellent results, whereas Gao et al. (2021) and Nam et al. (2019) maintained perfect values (figure 2D). Altogether, the external validation included 95 patients with 627 positive and 600 negative samples, highlighting the robust generalizability and performance of the models.

While direct benchmarking against existing tools is not possible due to its novel approach, baseline comparisons of hallmark-specific signature probabilities between normal (GTEx and GSE120795) and cancer datasets (TCGA, CCLE, POG570, PCAWG, TARGET, and MET500) reveal critical biological distinctions underlying tissue homeostasis and malignancy (figure 3, figure S6 appendix 1 pp 13-14). In normal datasets, hallmark activities exhibit tightly regulated density distributions, reflecting the stable, balanced biological processes necessary for maintaining normal cellular function. In contrast, cancer datasets display a marked shift, with significantly elevated probabilities for hallmark-specific signatures, indicating dysregulated processes that drive tumor progression. However, in certain datasets, there is variation in results, emphasizing that the model’s outcomes are inherently dependent on the quality and characteristics of the input data provided. The Kolmogorov-Smirnov (K-S) test further strengthened the findings with K-S statistic value above 0·7 with p-value of zero for all the hallmarks (table 2). These findings underscore the profound differences in hallmark activities between healthy and cancerous states.

**Figure 3:**
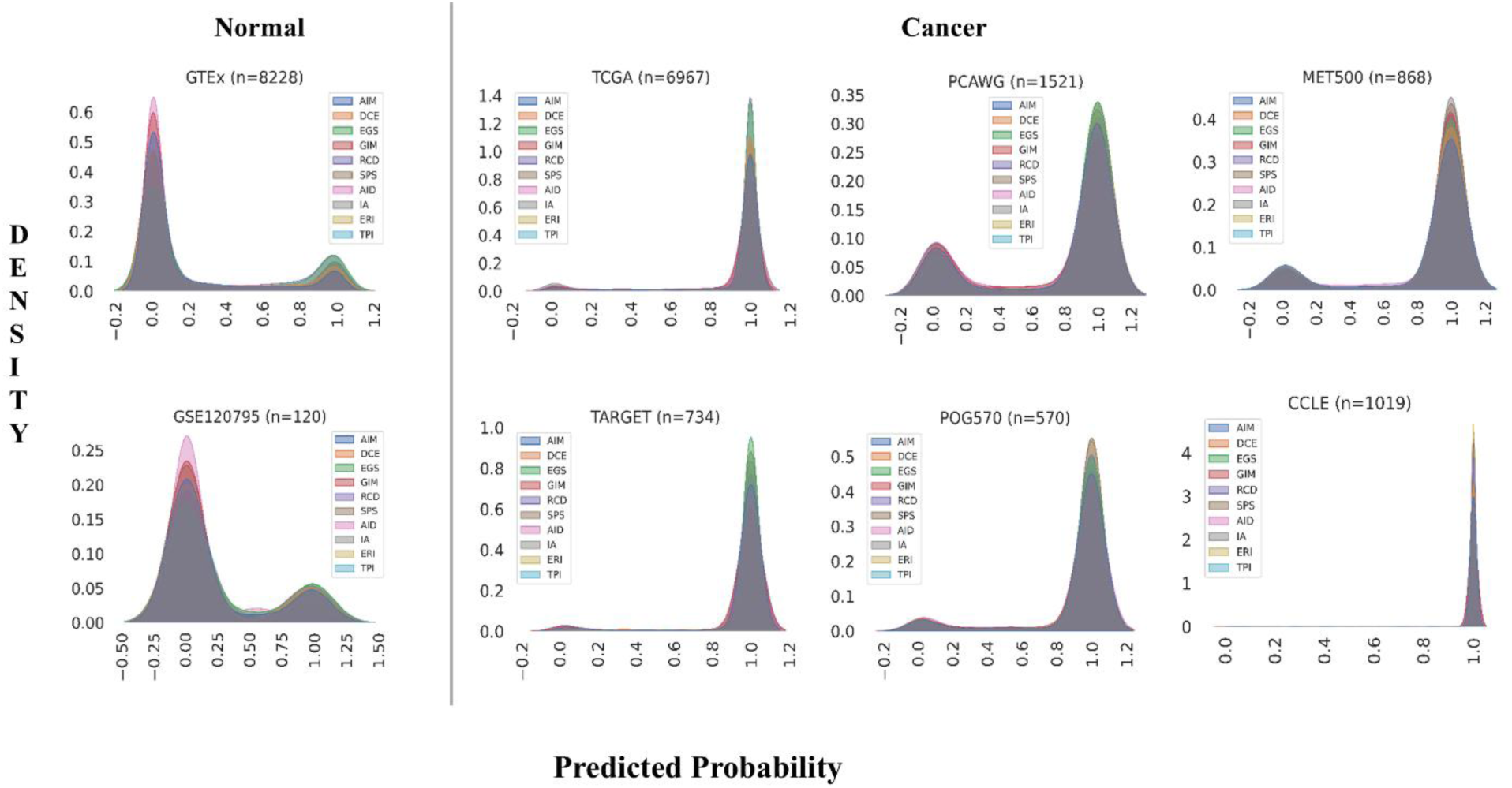
Probability distribution plots depicting predicted probabilities for hallmark-specific signatures are shown across normal and cancer datasets. The left panel includes normal datasets such as GTEx and GSE120795 (ANTE), showcasing distinct density distributions of predicted probabilities that emphasize hallmark-specific variations in normal tissues. The right panel features cancer datasets, including TCGA, CCLE, POG570, PCAWG, TARGET, and MET500, illustrating probability distributions that reveal a shift in hallmark activity between normal and cancer states. Each plot represents the predicted probabilities for ten hallmark-specific signatures.

**Table 2:** The Kolmogorov-Smirnov (K-S) test Statistic and p-value of the hallmark-specific probability difference in the model prediction.

|  | K-S Statistic | P-value |
| --- | --- | --- |
| AIM | 0.774614245 | 0 |
| DCE | 0.75721041 | 0 |
| EGS | 0.745627737 | 0 |
| GIM | 0.765449288 | 0 |
| RCD | 0.730226773 | 0 |
| SPS | 0.758095703 | 0 |
| AID | 0.768465534 | 0 |
| IA | 0.690902835 | 0 |
| ERI | 0.770428872 | 0 |
| TPI | 0.739581973 | 0 |

A detailed analysis reveals a dynamic progression of hallmark activity corresponding to different clinical cancer stages, offering valuable insights into the biological changes associated with tumor development and progression (figure 4). In the AJCC stages, hallmark activities progressively increase from Stage I to Stage IV, with the most significant co-association observed at advanced stages, underscoring the critical role of hallmark pathways in promoting tumor aggressiveness (figure 4A). A similar trend is observed in tumor stages (t1 to t4), where hallmark activity is highest in t4, reflecting the elevated engagement of these pathways in advanced and aggressive tumors (figure 4D). The metastasis stage (M0 to M1) reveals increased hallmark activity associated with metastatic potential, while the node stage (n0 to n2/3) demonstrates intensified activity with greater lymph node involvement, highlighting the role of hallmark pathways in tumor spread and metastasis (figures 4B and 4C). Collectively, these findings highlight the dynamic regulation of hallmark pathways during cancer progression and their potential as biomarkers for disease staging and therapeutic intervention.

**Figure 4:**
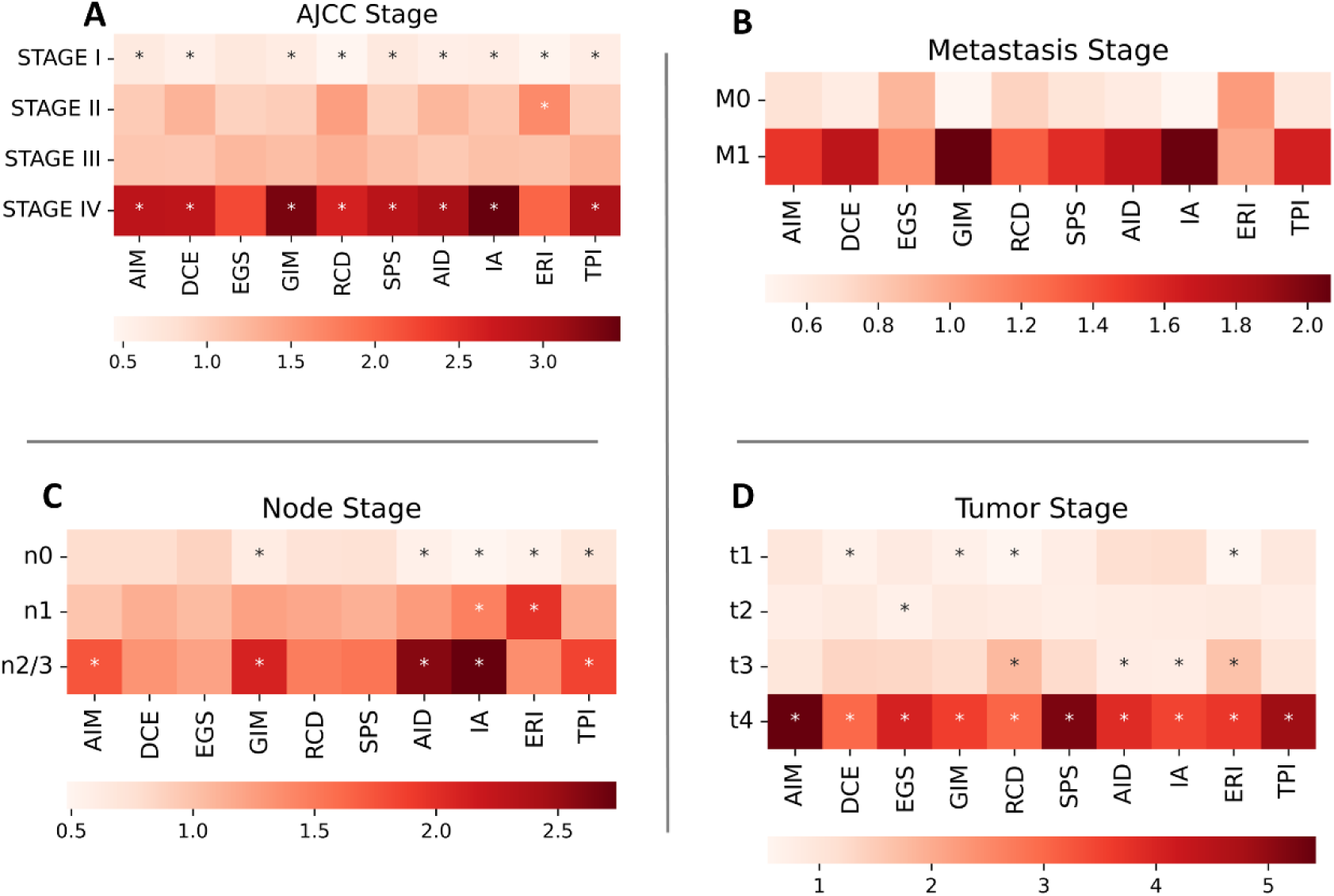
Heatmaps illustrate the association between hallmark activity and clinical cancer staging in TCGA datasets. A - AJCC stage, B - metastasis stage, C - node stage, and D - tumor stage. The color intensity in each heatmap reflects the magnitude of the association, with darker shades indicating higher associations. Asterisks (*) mark statistically significant associations (p-value < 0.05).

Further in-depth analysis highlighted the impact of hallmark-specific drugs on survival outcomes, shedding light on the intricate relationships between therapeutic interventions and cancer hallmark activities (figure 5). For Disease-Free Survival (DFS), hallmark ERI demonstrated strong associations with Anastrozole, AIM with Cyclophosphamide, TPI with Radiation, and RCD with Trastuzumab and Vinorelbine, as indicated by higher impact scores. These findings suggest that these therapies effectively target and modify the underlying hallmarks that drive tumor recurrence (figure 5A). In Progression-Free Survival (PFS), hallmark RCD was predominantly influenced by Cyclophosphamide, Trastuzumab, and Vinorelbine, reflecting these drugs’ potential to suppress disease progression. The high impact scores of these treatments highlight their efficacy in disrupting hallmark-specific pathways crucial for tumor growth and metastasis (figure 5B). For Overall Survival (OS), hallmark ERI was significantly modulated by Anastrozole and Leuprolide, RCD by Cyclophosphamide and Trastuzumab, GIM and IA by Irinotecan, and SPS by Radiation, as reflected by high impact scores (figure 5C). These results underscore the capacity of these therapies to enhance patient survival by effectively targeting hallmark pathways central to tumor progression.

**Figure 5:**
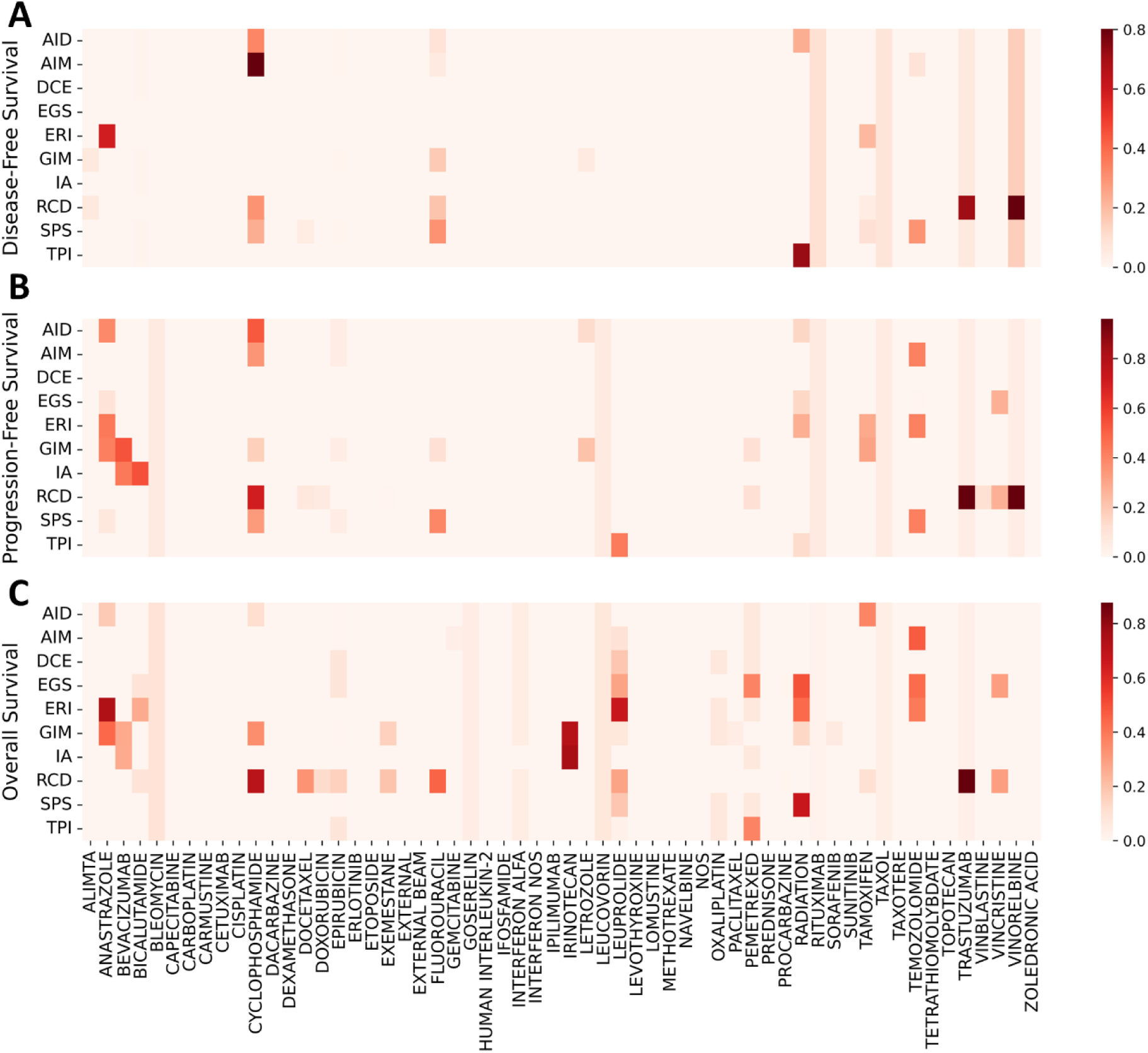
Impact of cancer drugs on Hallmarks of Cancer and Overall Patient Survival, Progression-Free Survival, and Disease-Free Survival. Color Intensity representing the effect of various drugs on distinct cancer hallmarks, with darker shades indicating higher impact scores, highlighting their contribution to improving patient survival outcomes.

## Discussion

Tumor staging and grading are indispensable for cancer assessment, offering insights into tumor size, spread, and cellular differentiation.^5^ However, these conventional metrics primarily reflect anatomical and morphological characteristics, often failing to capture the molecular mechanisms driving tumor behavior.^40^ Cancer progression is governed not only by physical growth but also by hallmark biological processes.^4^ Identifying and quantifying these hallmarks could provide a more nuanced understanding of tumor biology, uncovering therapeutic vulnerabilities and resistance mechanisms that traditional diagnostic methods overlook.^41,42^ Routine cancer diagnostics, such as imaging and histopathology, are limited in their ability to assess molecular hallmarks due to their focus on observable features rather than the underlying gene expression profiles that drive hallmark activation. Advances in transcriptomics and machine learning offer the potential to bridge this gap.^43^ By leveraging these tools, hallmark-specific molecular patterns can be identified and quantified, enabling more precise assessments of tumor biology. This approach may complement conventional methods, enhancing prognostic accuracy and supporting personalized therapeutic interventions.

OncoMark addresses these challenges by quantifying the activation states of cancer hallmarks using Multi-Task Neural Network trained on synthetic biopsy transcriptomics data. By identifying hallmarks, OncoMark enables stratification of tumors based on molecular profiles rather than solely anatomical characteristics. This molecular stratification may reveal biologically aggressive tumors that appear indolent through traditional grading and staging methods, improving risk assessment and early detection of relapse or resistance. Additionally, hallmark-driven profiling might uncover novel prognostic biomarkers, enhancing personalized care and disease outcome predictions.^44^ Furthermore, OncoMark capacity to capture hallmark interdependencies offers a comprehensive view of tumor heterogeneity, which is critical for advancing precision oncology.

Beyond prognosis, OncoMark offers significant potential in therapy design by identifying hallmark-specific vulnerabilities. For instance, tumors characterized by hallmark activation of angiogenesis may benefit from anti-angiogenic therapies, while those with genomic instability might respond to DNA damage repair inhibitors. OncoMark also ensures clinical relevance across diverse types of cancer, potentially supporting the development of tailored treatment strategies. Moreover, tracking hallmark states over time might help guide therapeutic adjustments and monitor treatment responses, which could contribute to improving patient outcomes.^42^

Despite its promise, the widespread adoption of OncoMark faces several challenges. The high cost of transcriptomic profiling and the need for specialized infrastructure limit accessibility in many healthcare systems, particularly in resource-limited settings. Integration into clinical workflows will require significant adaptations, including updates to diagnostic guidelines and training for healthcare providers. However, these challenges may diminish with technological advancements. The declining cost of sequencing, development of portable sequencing devices, and emergence of cloud-based analytical platforms could democratize access to transcriptomic analysis.^45^ Additionally, the growing integration of electronic health records and data-sharing initiatives may facilitate the incorporation of hallmark-based diagnostics into clinical practice.

In conclusion, OncoMark is, to our knowledge, the first computational tool to predict all hallmarks of cancer simultaneously. By bridging the gap between molecular biology and clinical practice, OncoMark has the potential to enhance prognostication, improve therapeutic targeting, and support the transition toward precision oncology. With continued advances in technology and infrastructure, hallmark-based diagnostics may become a routine component of personalized cancer management, offering patients more accurate and effective care.

## Contributors

SH and DG conceived the study. SP contributed to study and methodology design. SH and DG were responsible for overall project supervision. SP, CM and SB contributed to data collection. SP and CM prepared the initial manuscript draft. SP and CM contributed to figure design. SP and CM contributed to data analysis and result interpretation. DC provided invaluable insights throughout this study. BN cross-checked the study for accuracy and consistency. All authors contributed to the review of the manuscript.

## Data Sharing

All synthetic data generated in this study is publicly available at https://doi.org/10.5061/dryad.zw3r228jc. All codes generated in this study are publicly available at https://github.com/SML-CompBio/OncoMark. The web server can be accessed here: https://oncomark-ai.hf.space/. The python package is available here: https://pypi.org/project/OncoMark/. The comprehensive documentation to use OncoMark is available at https://oncomark.readthedocs.io/en/latest/. The single cell data used in this study is publicly available at https://www.weizmann.ac.il/sites/3CA/. MET500, TARGET, PCAWG data are publicly available at https://xenabrowser.net/datapages/. The TCGA data can be obtained from https://www.genome.gov/Funded-Programs-Projects/Cancer-Genome-Atlas, https://gdac.broadinstitute.org/, or https://www.cbioportal.org/. GTEx data is publicly available at https://www.gtexportal.org/home/. CCLE data is publicly available at https://sites.broadinstitute.org/ccle/datasets. ANTE data is publicly available at https://www.ncbi.nlm.nih.gov/geo/query/acc.cgi?acc=GSE120795. POG570 data is publicly available at bcgsc.ca/downloads/POG570/.

## Acknowledgments

We sincerely thank Amarnath Pal and Milan Patra for their valuable feedback on the initial manuscript draft, which helped us improve it. Our gratitude extends to the S.N. Bose National Centre for Basic Sciences and Ashoka University for their generous support and funding. S.H. acknowledges funding from the DST SERB Core Research Grant, while S.P, B.N, and D.G. are grateful to the Mphasis F1 Foundation for their support. The funders were not involved in the study’s design, data collection and analysis, decision to publish, or manuscript preparation. The views expressed in this article are those of the authors and do not necessarily reflect the policies, decisions, or perspectives of their employers or associated institutions. The results shown here are in whole or part based upon data generated by the TCGA Research Network (https://www.cancer.gov/tcga) and Therapeutically Applicable Research to Generate Effective Treatments (https://www.cancer.gov/ccg/research/genome-sequencing/target) initiative, phs000218.

## Correspondence

Correspondence should be addressed to Shubhasis Haldar and Debayan Gupta. Requests regarding codes and datasets should be directed to Shreyansh Priyadarshi.

## References

1 Lenz G, Onzi GR, Lenz LS, Buss JH, dos Santos JA, Begnini KR. The Origins of Phenotypic Heterogeneity in Cancer. Cancer Research 2022; 82: 3–11.

2 Swanton C, Bernard E, Abbosh C, et al. Embracing cancer complexity: Hallmarks of systemic disease. Cell 2024; 187: 1589–616.

3 Hanahan D, Weinberg RA. The Hallmarks of Cancer. Cell 2000; 100: 57–70.

4 Hanahan D, Weinberg RA. Hallmarks of Cancer: The Next Generation. Cell 2011; 144: 646–74.

5 Brierley J, O’Sullivan B, Asamura H, et al. Global Consultation on Cancer Staging: promoting consistent understanding and use. Nat Rev Clin Oncol 2019; 16: 763–71.

6 Greene FL, Sobin LH. The Staging of Cancer: A Retrospective and Prospective Appraisal. CA: A Cancer Journal for Clinicians 2008; 58: 180–90.

7 Bruni D, Angell HK, Galon J. The immune contexture and Immunoscore in cancer prognosis and therapeutic efficacy. Nat Rev Cancer 2020; 20: 662–80.

8 Su X, Lin Q, Liu B, et al. The Promising Role of Nanopore Sequencing in Cancer Diagnostics and Treatment. Cell Insight 2025;: 100229.

9 Granja JM, Klemm S, McGinnis LM, et al. Single-cell multiomic analysis identifies regulatory programs in mixed-phenotype acute leukemia. Nat Biotechnol 2019; 37: 1458–65.

10 Gao S, Tibiche C, Zou J, et al. Identification and Construction of Combinatory Cancer Hallmark–Based Gene Signature Sets to Predict Recurrence and Chemotherapy Benefit in Stage II Colorectal Cancer. JAMA Oncology 2016; 2: 37–45.

11 Dong R, Yang R, Zhan Y, et al. Single-Cell Characterization of Malignant Phenotypes and Developmental Trajectories of Adrenal Neuroblastoma. Cancer Cell 2020; 38: 716–733.e6.

12 Yost KE, Satpathy AT, Wells DK, et al. Clonal replacement of tumor-specific T cells following PD-1 blockade. Nat Med 2019; 25: 1251–9.

13 Pal B, Chen Y, Vaillant F, et al. A single-cell RNA expression atlas of normal, preneoplastic and tumorigenic states in the human breast. The EMBO Journal 2021; 40: e107333.

14 Gao R, Bai S, Henderson YC, et al. Delineating copy number and clonal substructure in human tumors from single-cell transcriptomes. Nat Biotechnol 2021; 39: 599–608.

15 Nam AS, Kim K-T, Chaligne R, et al. Somatic mutations and cell identity linked by Genotyping of Transcriptomes. Nature 2019; 571: 355–60.

16 Robinson DR, Wu Y-M, Lonigro RJ, et al. Integrative clinical genomics of metastatic cancer. Nature 2017; 548: 297–303.

17 Pleasance E, Titmuss E, Williamson L, et al. Pan-cancer analysis of advanced patient tumors reveals interactions between therapy and genomic landscapes. Nat Cancer 2020; 1: 452–68.

18 Ghandi M, Huang FW, Jané-Valbuena J, et al. Next-generation characterization of the Cancer Cell Line Encyclopedia. Nature 2019; 569: 503–8.

19 Aaltonen LA, Abascal F, Abeshouse A, et al. Pan-cancer analysis of whole genomes. Nature 2020; 578: 82–93.

20 THE GTEX CONSORTIUM. The GTEx Consortium atlas of genetic regulatory effects across human tissues. Science 2020; 369: 1318–30.

21 Suntsova M, Gaifullin N, Allina D, et al. Atlas of RNA sequencing profiles for normal human tissues. Sci Data 2019; 6: 36.

22 Luecken MD, Theis FJ. Current best practices in single-cell RNA-seq analysis: a tutorial. Molecular Systems Biology 2019; 15: e8746.

23 Heumos L, Schaar AC, Lance C, et al. Best practices for single-cell analysis across modalities. Nat Rev Genet 2023; 24: 550–72.

24 Menyhárt O, Harami-Papp H, Sukumar S, et al. Guidelines for the selection of functional assays to evaluate the hallmarks of cancer. Biochimica et Biophysica Acta (BBA) - Reviews on Cancer 2016; 1866: 300–19.

25 Zhang D, Huo D, Xie H, et al. CHG: A Systematically Integrated Database of Cancer Hallmark Genes. Front Genet 2020; 11. DOI:10.3389/fgene.2020.00029.

26 Iannuccelli M, Micarelli E, Surdo PL, et al. CancerGeneNet: linking driver genes to cancer hallmarks. Nucleic Acids Research 2020; 48: D416–21.

27 Liang P-I, Wang C-C, Cheng H-J, et al. Curation of cancer hallmark-based genes and pathways for in silico characterization of chemical carcinogenesis. Database 2020; 2020: baaa045.

28 Tate JG, Bamford S, Jubb HC, et al. COSMIC: the Catalogue Of Somatic Mutations In Cancer. Nucleic Acids Research 2019; 47: D941–7.

29 Menyhart O, Kothalawala WJ, Győrffy B. A gene set enrichment analysis for the cancer hallmarks. Journal of Pharmaceutical Analysis 2024;: 101065.

30 Cox DR. Regression Models and Life-Tables. Journal of the Royal Statistical Society: Series B (Methodological) 1972; 34: 187–202.

31 Andreatta M, Carmona SJ. UCell: Robust and scalable single-cell gene signature scoring. Computational and Structural Biotechnology Journal 2021; 19: 3796–8.

32 Murphy AE, Skene NG. A balanced measure shows superior performance of pseudobulk methods in single-cell RNA-sequencing analysis. Nat Commun 2022; 13: 7851.

33 Crowell HL, Soneson C, Germain P-L, et al. muscat detects subpopulation-specific state transitions from multi-sample multi-condition single-cell transcriptomics data. Nat Commun 2020; 11: 6077.

34 Qiu X, Wu H, Hu R. The impact of quantile and rank normalization procedures on the testing power of gene differential expression analysis. BMC Bioinformatics 2013; 14: 124.

35 Ioffe S, Szegedy C. Batch Normalization: Accelerating Deep Network Training by Reducing Internal Covariate Shift. In: Proceedings of the 32nd International Conference on Machine Learning. PMLR, 2015: 448–56.

36 Caruana R. Multitask Learning. Machine Learning 1997; 28: 41–75.

37 Smirnov N. Table for Estimating the Goodness of Fit of Empirical Distributions. Ann Math Statist 1948; 19: 279–81.

38 Shiryayev AN. 15. On The Empirical Determination of A Distribution Law. In: Shiryayev AN, ed. Selected Works of A. N. Kolmogorov. Dordrecht: Springer Netherlands, 1992: 139–46.

39 Cornfield J. A method of estimating comparative rates from clinical data; applications to cancer of the lung, breast, and cervix. J Natl Cancer Inst 1951; 11: 1269–75.

40 Bello DM, Russell C, McCullough D, Tierno M, Morrow M. Lymph Node Status in Breast Cancer Does Not Predict Tumor Biology. Ann Surg Oncol 2018; 25: 2884–9.

41 Hanahan D. Hallmarks of Cancer: A 2012 Perspective. Annals of Oncology 2012; 23: ix23.

42 Bailón-Moscoso N, Romero-Benavides JC, Ostrosky-Wegman P. Development of anticancer drugs based on the hallmarks of tumor cells. Tumor Biol 2014; 35: 3981–95.

43 Wei L, Niraula D, Gates EDH, et al. Artificial intelligence (AI) and machine learning (ML) in precision oncology: a review on enhancing discoverability through multiomics integration. British Journal of Radiology 2023; 96: 20230211.

44 Zhou Y, Tao L, Qiu J, et al. Tumor biomarkers for diagnosis, prognosis and targeted therapy. Sig Transduct Target Ther 2024; 9: 1–86.

45 Loose M, Malla S, Stout M. Real-time selective sequencing using nanopore technology. Nat Methods 2016; 13: 751–4.

